# Cognition and brain gene expression vary with sex and body size in a polyphenic dung beetle (*Onthophagus taurus*)

**DOI:** 10.64898/2026.10.14.759186

**Authors:** R. R. Westwick, A. P. Moczek

## Abstract

Individual variation in cognition can have critical fitness consequences for animals. Therefore, it is important to understand the relationship between cognition and ecologically relevant life history traits that vary within species. Furthermore, we lack a unified understanding of how these traits are integrated mechanistically with the cognitive outcomes they are associated with. We examined the relationship between cognitive performance on multiple tests on one side—and sex, a nutritionally-cued reproductive male polyphenism, and body size on the other—as well as their respective relationships to brain gene expression. Using the bull-headed dung beetle (*Onthophagus taurus*), we performed an associative learning assay, a spatial cognition assay, and a detour test on large fighter males, small sneaker males, and correspondingly large and small females. We then performed RNAseq on the brains of the beetles that had performed these assays. We found that sex was an important predictor of performance on the spatial cognition assay, while body size (which is closely related to early life nutrition) was associated with performance on the detour task. The associative learning assay did not yield significant results. RNAseq data revealed complex and multifaceted relationships between brain gene expression and our factors of interest. This work provides insights into the complex links between cognition, brains, and ecologically relevant traits.

## BACKGROUND

As animals move about the world, they must make decisions about survival and reproduction, such as where and how to forage or which mate to choose. The decisions that they make can have important consequences for their fitness; for example, parasitoid wasps that are allowed to learn which substrates are more likely to have viable host eggs to parasitize are able to produce more offspring than wasps that are not allowed to learn (Dukas & Duan, 2000). Cognition—the ways in which animals take in information through the senses, process, retain and decide to act on it—thus serves as an important intermediate between the informational input received from an organism’s environment and the functional fitness consequences of its behavioral response to that information (Dukas, 2004; Shettleworth, 2001). Consequently, individual differences in cognition often lead to differential fitness effects between these individuals (Boogert et al., 2018).

Given the importance of individual cognitive differences, it is important to understand what factors underlie them. Influences that are known to be associated with individual variation in cognition include intrinsic (e.g. neurobiological correlates, genetics) and extrinsic (e.g. the physical environment, the social environment) factors (Ashton et al., 2018; Croston et al., 2015; Dubois et al., 2018; Sauce et al., 2018). Yet many other life history features exist that feed into an organism’s biological state. In particular, how features such as early life nutrition, alternative male reproductive tactics, or sex relate to the spectrum of individual cognitive variation within a species is less well understood, let alone how these features act and interact within a whole organism.

One important mechanism that could link functional traits to cognition across the whole organism is gene expression. Even though cognition is primarily understood as an emergent property of complex neuronal processes, these processes are themselves underlain by tissue- and cellular-level properties, actions, and responses, all of which in turn necessitate context-specific gene expression. Yet how cognitive performance is related to brain gene expression, and how that relationship is shaped by factors such as sex, male reproductive morph, or early life nutrition, remains poorly understood.

To investigate how key functional traits are related to cognition and brain gene expression, we used the bull-headed dung beetle, *Onthophagus taurus. O. taurus* is a dung beetle that feeds primarily on ungulate dung and spends much of its multi-month adult life excavating subterranean tunnels underneath dung pads, flying to new dung patches every few days and assessing patch quality while foraging (Dormont et al., 2004; Hanski & Cambefort, 2014). These insects make an excellent model for testing the relationship between cognition, brain gene expression, and functional traits. Male and female *O. taurus* exhibit striking sexual dimorphism in both morphology and behavior (Fig. 1a&b). Additionally, within males there is a nutritionally-cued male polyphenism in morphology and reproductive tactic (Fig. 1a&c; (Moczek & Emlen, 2000)).

**Figure 1:**
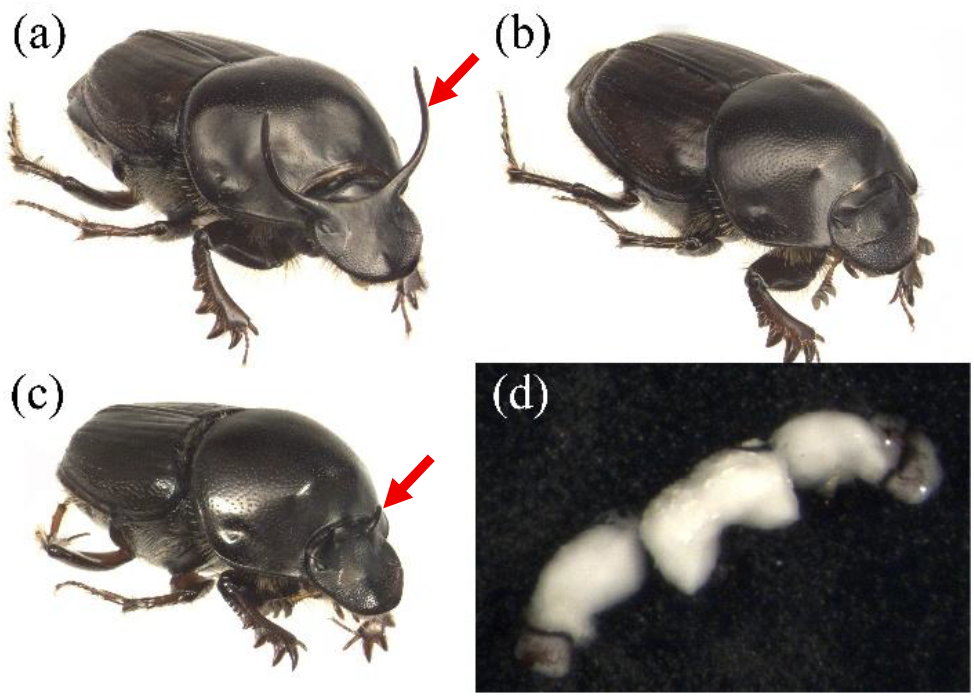
*Onthophagus taurus* show striking sexual dimorphism and male reproductive polyphenism. a) “Major” fighter male with large head horns (shown with red arrow); b) female; c) “minor” sneaker male with small head horns (shown with red arrow); d) transverse view of a dissected *O. taurus* brain (after flash-freezing and lyophilization); optic lobes are the dark regions on the far lateral edges, antennal lobes are the paired protrusions on the anterior edge. (*Beetle photos provided courtesy of Erica Nadolski. Brain photo taken by Rebecca Westwick*.)

Males who receive high nutrition in early life develop into “major males” with large bodies and exaggerated head horns, and who primarily use fighting behavior to aggressively defend tunnels with females against other males (Fig. 1a). Males who receive lower nutrition in early life develop into “minor males” with smaller bodies and rudimentary head horns (Fig. 1c; (Moczek, 1998)). Minor males will also fight other males to monopolize tunnels but will switch to sneaking tactics if fighting fails; major males never perform these sneaking behaviors. These sneaking behaviors can be diverse, including behaviors such as waiting for the tunnel-defending major male to be distracted in a fight before sneaking past him, digging side intercept tunnels to access the defended tunnel below where the defending male is positioned, and securing matings up at the dung pad (Moczek & Emlen, 2000). Females show a similar range of body sizes as males based on early life nutrition, but their behavioral repertoire is consistent across their body size range (Fig. 1b). In particular, females show complex parental care behaviors (sometimes but not always assisted by males), constructing specialized “brood ball” nurseries for each egg they lay and modifying their care behaviors in response to environmental triggers such as dung quality, temperature, and heavy metal pollution ((Hunt & Simmons, 2004; Hunt et al., 2002; Macagno et al., 2018; Moczek, 1999), Towse et al. *forthcoming*). Thus, *O. taurus* offers a promising opportunity to compare the effects of sex, reproductive tactic, and early life nutrition (which is strongly correlated with body size) on cognition and brain gene expression.

We performed a battery of cognition tests (an associative learning assay, a spatial cognition assay, and a detour task) on a suite of beetles (large major males, small minor males, and correspondingly large and small females). We then performed RNAseq on the brains of the beetles that had completed these cognition assays to assess baseline brain gene expression in these individuals.

We expect all of our functional groups to use associative learning and spatial cognition in nature while assessing dung quality and navigating their subterranean tunnels respectively; we therefore hypothesized that the associative learning and spatial cognition assays would reveal cognitive variation between individuals, but that performance would be similar on average across our functional groups. Conversely, for the detour task, we hypothesized that minor males—who have to facultatively switch between fighting and sneaking tactics in reproductive contests, including sneaking behaviors that involve physically approaching a goal from an alternative angle—would outperform the other functional groups. Contrary to our predictions, we found that sex was an important predictor of performance on the spatial cognition assay (with males outperforming females) and that body size—a proxy for early life nutrition—was an important predictor of performance on the detour task (with large individuals outperforming small individuals). We additionally hypothesized that brain gene expression would (i) covary in part across functional groups, thereby revealing both shared and distinct gene expression repertoires (ii) covary as a function of cognitive performance; and (iii) reveal diverse genes with statistical interaction effects between our factors of interest. All of these brain gene expression hypotheses were supported by our data.

Overall, this work provides an important line of evidence that ecologically relevant traits such as sex and body size (i.e. early life nutrition) are associated with differences in cognitive performance within a species. This work additionally uncovers brain gene expression profiles associated with these traits, providing evidence for potential genetic and physiological mechanisms underlying the cognitive differences established in this work.

## METHODS

### Beetle Husbandry

*Onthophagus taurus* beetles used in this experiment were sourced from a large lab colony that were descendants (generations F1+) of haphazardly mixed-mated individuals wild-caught in North Carolina, USA. The source colony population remained between 100-200 individuals throughout the experiment, even as individuals were removed for this experiment, with new individuals being routinely added that were offspring from breeding groups of the F0 wild-caught and F1 lab-reared generations. The colony was kept in a large plastic container with approximately five inches of a damp sand-soil mixture, with a pad of lightly homogenized cow dung on window screen mesh being added twice per week as a food source. The colony was kept in a climate-controlled room at approximately 24°C and 25% relative humidity with a 16h light/8h dark photoperiodic cycle, an approach our lab has used to successfully rear and breed these beetles for decades (Schwab & Moczek, 2016; Shafiei et al., 2001).

We haphazardly sampled experimental beetles (N = 48, 12 each of major males, minor males, large females, and small females) from the lab-reared colony two days before that beetle’s first cognition assays. Up to 8 beetles were in progress at a time. Individuals were collected, sexed, had their thorax width measured (to determine body size, which is also a strong proxy for early life nutrition (Moczek, 1998)), and (for males) were recorded as being major or minor based on horn length. From then on, beetles were kept in small microcosms, each with two same-sex beetles. While males will fight with one another if females are present, we have previously observed that same-sex individuals will typically not interact with one another if the other sex is *not* present. Males will also exhibit courtship and guarding behaviors of females if they are kept together. Therefore, keeping individuals in same-sex pairs allowed us to decrease the housing burden that would come with keeping every beetle alone, while minimizing the potential disruptions that arise when mixed-sex groups are housed together in small spaces. The microcosms contained approximately one inch of the same lightly damp sand-soil mixture, a small piece of window screen mesh, and a small dung pad sufficient to support two beetles.

We removed food for a period of approximately 48 hours prior to each assay as a means to ensure the beetles would be motivated by a food reward. *Onthophagus taurus* are typically quite robust to starvation; we do not see excess mortality until around a week (or even longer) following food removal in the absence of other stressors, so this two-day deprivation is expected to make them hungry but not cause harm. We did not observe excess mortality during this experiment as a result of the food deprivation (beetles were sourced from a mixed-age colony, so a small amount of mortality was documented during the experiment as we did not precisely know the ages of the beetles when they were sourced; but this mortality was not elevated relative to null expectations within the colony).

### Cognition assays

#### Assay schedule

All rounds of each assay type occurred on a single day. The assays were performed in a separate climate-controlled room from where the main colonies were kept, but the climatic and light conditions were identical. Assays were typically completed within the time range of 09h00 and 17h00. We randomized whether the associative learning or spatial cognition test was performed first, while the detour test was always performed last of the three. We did this to ensure that the beetles had multiple experiences with the novel food reward used (*see below*), since the detour test required them to detect it from afar. In between the different assay types, the beetles were given 48 hours to feed, then underwent another 2-day food deprivation before the next assay. Thus, a complete schedule for a single beetle would consist of: sourcing and measuring, a 2-day food deprivation, the first test day (associative or spatial), a 2-day feeding period, a 2-day food deprivation, the second test day (the other of either associative or spatial), a 2-day feeding period, a 2-day food deprivation, the detour test day, then a 2-to-4 day feeding period before euthanasia via flash-freezing for brain collection.

#### Rewards

While larval *O. taurus* chew and eat the fibrous dung components, adults are filter feeders and consume only the liquid components of the dung and tiny particles dissolved therein. We therefore leveraged this liquid content as a way to better standardize the food reward used in all assays. We first took approximately 2 kg of frozen-and-defrosted dung collected from grass-fed beef cattle at a near-organic farm (Marble Hill Farm, Bloomington, IN, USA). We then squeezed the dung thoroughly through a cheese cloth, saving the extruded liquid that we will hereafter call “dung tea.” Dung tea was aliquoted into small containers and re-frozen, such that the amount for a single day’s assays could be thawed at a time. We created a set of one-inch sponge cubes that we soaked in the dung tea that were presented as the reward during assays, refreshing the liquid every 30 min – 1 hr to keep it fresh and damp. Early pilot results showed that beetles given a dung tea sponge in a large open arena showed strong attraction to the sponge, spending around 90% of their time in contact with it and drinking the liquid from the sponge.

#### Associative learning assay

For the associative learning assay, we tested color learning. We created an arena out of white LEGO™ bricks with a large starting chamber and two smaller chambers side-by-side on the same wall (Fig. S1a). Each chamber had a large entry door constructed of different colored LEGOs™—one door was light green, the other was dark blue. Color vision has not been functionally tested in this species, but they are known at a minimum to have genes that code for opsins that sense green and UV light, and may have trichromatic vision (as many insects do) due to an opsin duplication restoring blue sensitivity (Sharkey et al., 2017). Therefore, we expect that these colors would be within their range of spectral sensitivity. Furthermore, the contrast of the lighter green versus the darker blue against the white background would be expected to enhance their ability to distinguish between the two doors. Each beetle was randomly selected to have the green or blue door be the rewarding color. The beetles were introduced into the starting chamber under a dark acclimation cup and allowed to rest for 5 minutes before every round. The first round (a “Round 0”) was a no-choice round, where only the “correct” door was in place and the other chamber was blocked by solid white wall. Every subsequent round included both doors. Both chambers contained a dung tea sponge reward to release scent cues, but the dung tea sponge in the “incorrect” chamber was blocked by a permeable barrier that allowed airflow but did not allow the beetle to contact the reward. We randomized whether the “correct” side would be on the left or the right for each round, and the LEGO™ design allowed us to easily swap which door was on which side. Following the no-choice round, the beetles were given 10, 15-minute training rounds followed by a final 15-minute test round. A live observer recorded the time it took for the beetle to choose a chamber to enter and the time it took for the beetle to reach the reward. The trial was stopped after one of three conditions: 1) if the beetle fully entered the correct chamber and made contact with the “correct” reward, at which point the beetle would be allowed to interact with the reward for 2 minutes; 2) if the beetle fully entered the incorrect chamber made contact with the barrier blocking the decoy inaccessible “incorrect” reward, at which point the trial would be stopped immediately and the beetle removed; or if 15 minutes elapsed and no choice was made. The arena was thoroughly cleaned with 70% ethanol and deionized water between every round.

#### Spatial cognition assay

For the spatial cognition assay, we created a small four-chamber maze out of black LEGO™ bricks. The maze design was functionally a double-T maze (Fig. S1b); after a five-minute acclimation period, the beetle was released from an acclimation chamber at the bottom of the arena and allowed to walk forward. There the beetle encountered the first T-junction, a long horizontal center hallway in which the beetle needed to choose to turn right or left. This hallway dead-ended on both sides into a second set of T-junctions, where the beetle once more had to choose to turn either right or left. The beetle would then encounter one of four chambers. One chamber was “correct” and contained an accessible reward. The other three chambers were “incorrect” and contained decoy rewards that released the same scent cues, but were blocked by small doors with holes that allowed airflow but did not permit the beetle to enter and interact with the reward. Each beetle’s “correct” chamber (top left, top right, bottom left, bottom right) was randomized for that individual beetle, but was kept the same throughout all rounds. These trials were conducted with a red cellophane lid over the top of the maze. As our beetles are not believed to have photoreceptors that detect the color red, this setup would more closely simulate the dark, subterranean tunnels that the beetles navigate in nature (and in their home cages) while still allowing human observation. The beetles were given 8, 15-minute rounds to learn the location of the correct chamber. A live observer recorded how long it took the beetle to leave the acclimation area, how long it took them to make their first choice, whether that first choice was correct or not, how long it took them to find the correct chamber (if not on the first try), and how many incorrect choices were made before reaching the correct chamber. The trial was stopped after the beetle had reached the correct chamber and interacted with the reward for 2 minutes, or if 15 minutes had passed and the beetle failed to reach the correct chamber. The arena was thoroughly cleaned with 70% ethanol and deionized water between every round.

#### Detour task

For the detour task, we used a 36 cm x 18 cm plastic arena. A grid was drawn on the floor to aid tracking of the individual’s movements. We constructed two identical barriers from LEGO™ bricks and either window screen mesh (test apparatus) or flexible black polyethylene plastic (control apparatus). From above, this barrier appears as an upside-down U that is 6.4 cm on each long side, 4.8 cm on the short side, and is 5.6 cm tall (Fig. S1c). The beetle was placed in a small, dark acclimation chamber inside of this barrier, with an accessible reward on the opposite side of the barrier and the open face of the “U” facing away from that reward. Thus, if the beetle initially moved towards the reward, it would encounter the barrier and then be forced to figure out how to maneuver around it to get to the reward. The beetles were given one control round to start. During the control round, the apparatus with the (impermeable) black plastic was used. The beetle would be given a 5-minute acclimation period under a dark acclimation cup. Then, when the acclimation cup was lifted, the beetle would be free to roam around the arena. A live observer recorded how long it took the beetle to contact the barrier, how long it took the beetle to leave the area within the barrier, where the beetle went within the arena (with the help of the grid on the floor of the arena), and how long it took the beetle to contact the reward. The trial would be stopped when the beetle contacted the reward, at which point it would be given two minutes to interact with the reward, or when 30 minutes had passed without the beetle successfully making contact with the reward. The test rounds were run similarly, but with the mesh barrier (that the beetle can presumably see/smell the reward through) rather than the impermeable plastic barrier. Each beetle received only one control round and one test round, as we wanted to capture the initial problem solving of getting around the barrier rather than a learned route that would develop over multiple rounds with the test mesh apparatus. The arena and apparatuses were thoroughly cleaned with 70% ethanol and deionized water between every round.

### RNAseq

#### Freezing and tissue preservation

We did not enter this experiment with an *a priori* expectation about which assay would be most salient for our brain gene expression results. We therefore elected to capture baseline brain gene expression, rather than pairing our brain collection with any assay in particular. Thus, after the final assay (the detour test), the beetles were given 2-4 days in their home microcosm with food *ad libitum* to return to a neutral state. We then flash-froze the beetles in a slurry of dry ice and ethanol and preserved them in microfuge tubes at -80°C until brain dissection.

#### Dissections

All surfaces and tools were thoroughly cleaned with ethanol and RNAse AWAY™ throughout the dissection and RNA extraction processes. We first removed the beetles’ heads using a sharp scalpel (Ambler Surgical, 7516) on a silicone-lined dissection dish (Pyrex, DD-90-S-BLK) that was pre-chilled on dry ice to maintain low temperature. We then transferred each head into an individual well of a pre-chilled 12-well plate (GET SPECS) that was covered with a Kimwipe (Kimtech, 34155) and secured with a rubber band. We then partially lyophilized the frozen heads in a Labconco FreeZone 6L Console Freeze Dryer at 50 Pa, -48°C for 45 minutes to help separate the different tissue types with the head capsule. The lyophilized heads were then returned to the chilled dissection dish with a thin layer of 100% ethanol (which helps float away any extraneous tissues), everything continuing to sit on a bed of dry ice to maintain temperature. The dissection dish setup was then viewed under a stereo microscope (Leica, MZ16). Using the scalpel and two sets of 5.SA ultrafine tweezers, we carefully chipped away pieces of cuticle and popped off the eye caps. We then scraped away muscle, glandular tissue, and connective tissue until the brain was uncovered. We used the flat of the scalpel to carefully lift the brain out of the head capsule and isolate it, inspecting it to ensure complete removal of all extraneous tissues (Fig. 1d). We transferred the isolated whole brain to a pre-chilled microfuge tube with 300 uL of TRIzol™ reagent and aided lysis by mechanically disturbing the tissue with a sterilized pestle. The tube of TRIzol™ with tissue was stored at -80°C until RNA extraction.

#### RNA extraction and sequencing

We extracted RNA using a Direct-zol RNA MiniPrep w/TriReagent kit (Zymo, R2053-A) according to the manufacturer’s instructions. RNA quantity and quality were measured on a NanoDrop microvolume spectrophotometer (Thermo Fisher, 13-400-525) and an Agilent 2200 TapeStation. One sample failed quality metrics; therefore, though the sample size for the behavioral assays is N = 48, the sample size for the brain RNA expression is N = 47 (11 major males, 12 minor males, 12 large females, 12 small females). Library preparation and paired-end sequencing were carried out by the Indiana University Center for Genomics and Bioinformatics using an Illumina TruSeq Stranded mRNA High-Throughput kit and NextSeq 2000 (Illumina, 20038897) using a 50-cycle P3 flow cell. Average sequencing depth was 34.6 million reads per sample and average sample Mean Quality Score was 38.98/40.

### Data Analysis

#### Data analysis—behavioral data

We assessed both the associative learning assays (measuring “correct” vs “incorrect”, excluding rounds where an individual did not make a choice, which were rare) and the spatial cognition assays (measuring if the first chamber the beetle approached in a given round was “correct” or “incorrect”, excluding rounds where an individual did not make a choice, which were rare) using generalized linear mixed models. We used the glmer::lme4 function with sex, body size, and round as fixed effects and individual ID as a random effect; our models were run as binomial family with a logit link (Bates et al., 2015). Model diagnostics were assessed with the “DHARMa” package, which includes a QQ plot, Kolmogorov-Smrinov test, a dispersion test, an outlier test, a within-group uniformity test, and a Levene test (Hartig, 2022). For the spatial cognition assay, we additionally ran a binomial test (binom.test::stats) on the final round separately for males and females to test performance versus chance, using 0.25 threshold due to the 4-chamber design (R Core Team, 2021). For the detour test, due to its nature of not having a binary “correct” versus “incorrect” choice and truncation of an event that would almost certainly happen eventually if given infinite time, we analyzed with a Cox Proportional Hazard Model using packages “survival” and “survminer” (Kassambara et al., 2026; Therneau, 2020). Plots were created using “ggplot” and “viridis,” as well as built-in functions from previously mentioned packages (Garnier et al., 2021; Wickam, 2016).

#### Data analysis—RNAseq data

We used FastQC (version 0.11.9) and Trimmomatic (version 0.36) to perform quality control and trim adapters (Andrews, 2010; Beier et al., 2026). These reads were then fed into Salmon (version 1.10.1) to generate abundance estimates by pseudoaligning reads to a published *Onthophagus taurus* transcriptome (Davidson & Moczek, 2024; Patro et al., 2017). All data analyses were carried out in R version 4.6.1 (R Core Team, 2021). We used the *tximport* function to import abundance estimates and convert to transcript counts (Soneson et al., 2015).

To facilitate the inclusion of cognitive performance in the transcriptomic analysis, which requires a categorical variable, we created a composite measure of cognitive performance. This composite measure was created by splitting the top-performing beetles from the bottom-performing beetles *within each functional group* (e.g. the top 6 large females were “high” performers while the bottom 6 large females were considered “low” performers; similar for small females and major and minor males). These groups were primarily based on the success rate on the final rounds of the spatial cognition and detour assays (the associative learning assay was excluded from the composite measure due to low overall success rates, see “Results”). After success rate, we used a series of tie breaks to rank-order the beetles within each functional group: spatial efficiency at solving the detour test, the number of incorrect choices before the correct solution on the spatial cognition assay, then the average latency to solve the spatial cognition and detour assays. Once the beetles were in rank order, the top 6 within each functional group were marked as “high” performers and the bottom 6 were marked as “low” performers.

To remove genes with low read counts, we removed genes that did not meet the criteria of having at least 10 counts-per-million in at least 5 samples. We then used the “DESeq2” package to create a DESeq2 dataset using a design = ∼Sex*Size*Performance and specified all contrasts to incorporate sex, body size (large versus small), a composite measure of cognitive performance (high versus low, described above), and all two- and three-way interactions (Love et al., 2014). We used the DESeq2::lfcShrink function with the “ashr” type specification to shrink log2 fold change estimates for genes with high dispersion (Stephens, 2017). To assess differentially expressed genes, we included genes with a shrunken log2 fold change of at least 0.4 and a Benjamini-Hochberg-adjusted p-value of under 0.05 (Benjamini & Hochberg, 1995). We used the “ggplot2” and “viridis” packages for plot generation (Garnier et al., 2021; Wickam, 2016). Packages such as “tidyr” and “stringr” were used to facilitate data manipulation (Wickham, 2025; Wickham et al., 2026). Plots were arranged and plot labels were modified (e.g. to change font and text size) with Adobe Illustrator.

## RESULTS

### Cognition Assays

#### Associative Learning

Our associative learning assay did not yield results to indicate that the beetles were learning the association, nor were there significant differences between our groups (generalized linear mixed model: ANOVA: *Round*: Wald X^2^_1_=0.19, p= 0.66; *Sex*: Wald X^2^_1_=0.05, p= 0.82; *Size*: Wald X^2^_1_=2.29, p= 0.13; *Sex*Size*: Wald X^2^_1_=0.54, p= 0.46). This apparent lack of learning could have been due to the small number of training rounds (11 rounds). A small pilot study had indicated that this length of training would be sufficient, as learning was demonstrated in only 8 rounds. But there were key differences between the pilot study and our main data collection: namely, the pilot study was conducted with wild-caught individuals, while the main data collection was carried out with lab-reared individuals. Future experiments could explicitly test the role of rearing environment on associative color learning in this species.

#### Spatial Cognition

For our spatial cognition assay, we found evidence that the beetles were able to learn the correct route within the 8 training rounds. This result was not consistent across all groups, however. We found that males and females showed significant differences in performance throughout the training rounds, with males outperforming females (Fig. 2; generalized linear mixed model: ANOVA: *Round*: Wald X^2^ =5.50, p= 0.02; *Sex*: Wald X^2^ =13.69, p= 0.0002; *Size*: Wald X^2^ =2.32, p= 0.13; *Sex*Size*: Wald X^2^ =2.63, p= 0.10). During the final round in particular, males performed significantly better than chance, while females did not (binomial test: *Males:* p= 0.008; *Females:* p= 1). In contrast, body size was not associated with differences in performance on this assay.

**Figure 2:**
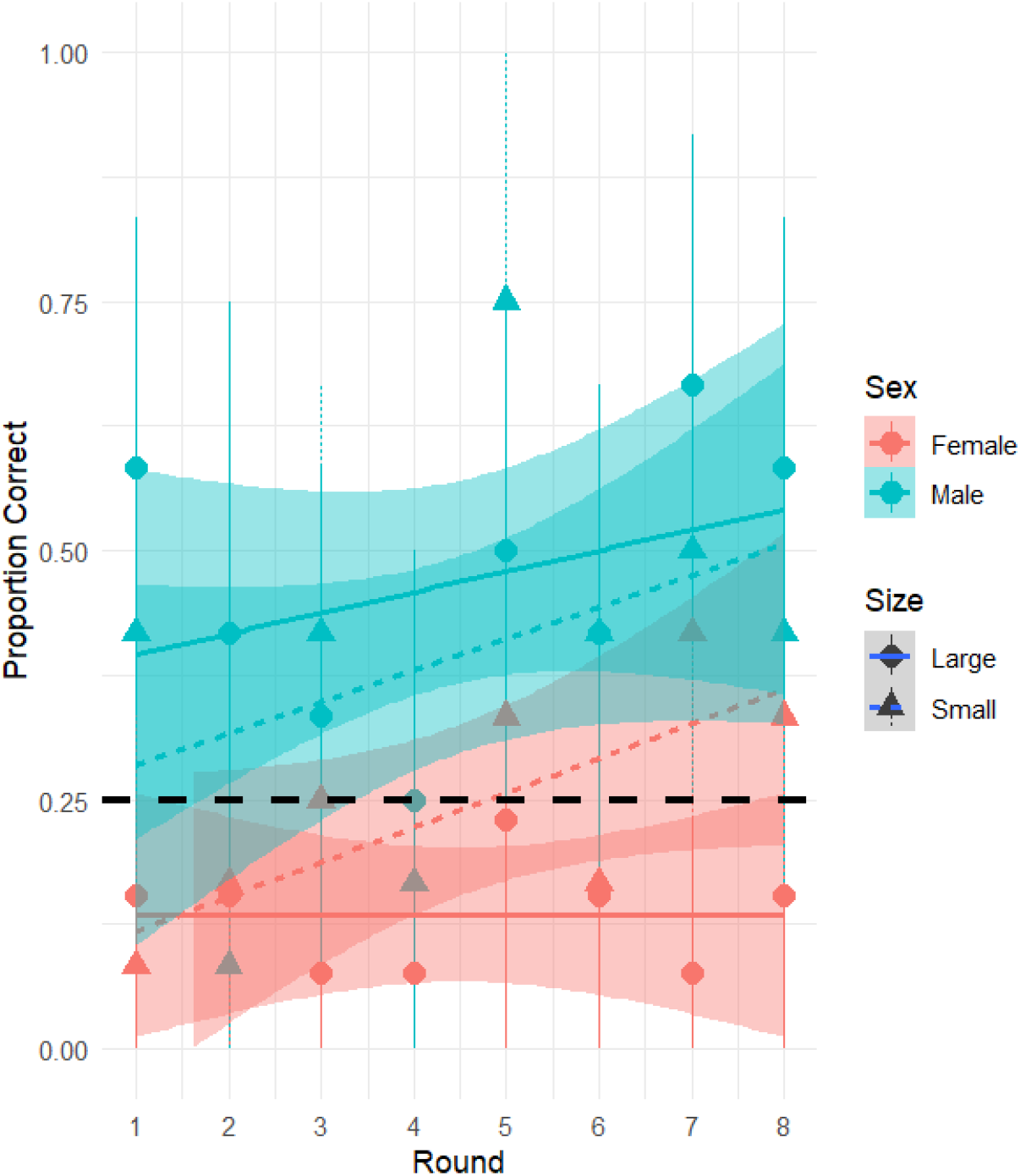
Male beetles outperformed female beetles (regardless of body size) on successive training rounds of a spatial cognition assay. Dotted black line represents the random chance threshold (0.25). Male data points are in blue, female in red, with large data points in circles with solid trendlines and small data points in triangles with small-dashed trendlines. Points with bars show the mean and 95% bootstrapped confidence intervals for each subgroup at each round, while the trend lines are linear regressions for each subgroup.

#### Detour Task

First, to validate our experimental approach for our detour test, we examined how the beetles responded to the control apparatus versus the test apparatus. We observed that the beetles almost universally approached the apparatus in the direction of the reward upon the beginning of the trial rather than walking out of the open side of the apparatus on the opposite side as the reward (75/76 trials). We found that (as expected), beetles spent longer attempting to approach the reward through the permeable mesh than the impermeable plastic (i.e. the beetles left the barrier sooner and maneuvered around it for the control apparatus; paired t-test: *Control*: mean = 203.8, SD = 392.4; *Test*: mean = 677.3, SD = 652.7; *t*_*37*_ = -3.63, *p* = .0004).

For our previous assays, we found that using body size as a categorical variable produced the best models as assessed by AIC. This designation of categorical body size (large vs small) is reflective of the male reproductive polyphenism. In contrast, for our detour test, we found that model performance was superior when we instead used a continuous measure of body size (pronotum width); despite the sharp cutoff for the distinct male “major vs minor” polyphenism in head horn morphology and reproductive behavior, body size itself varies more or less continuously underneath this division (Moczek, 1998).

When testing our groups of interest, we found that large individuals trended towards outperforming small individuals, regardless of sex (Fig. 3, note that we separate by “large vs small” categorically for visualization purposes, but the statistical model was run with “pronotum width” as a continuous measure of body size; Cox proportional hazard model: Z=1.70, p=0.088). In contrast to our spatial learning assay, sex was not associated with differences in performance on the detour test.

**Figure 3:**
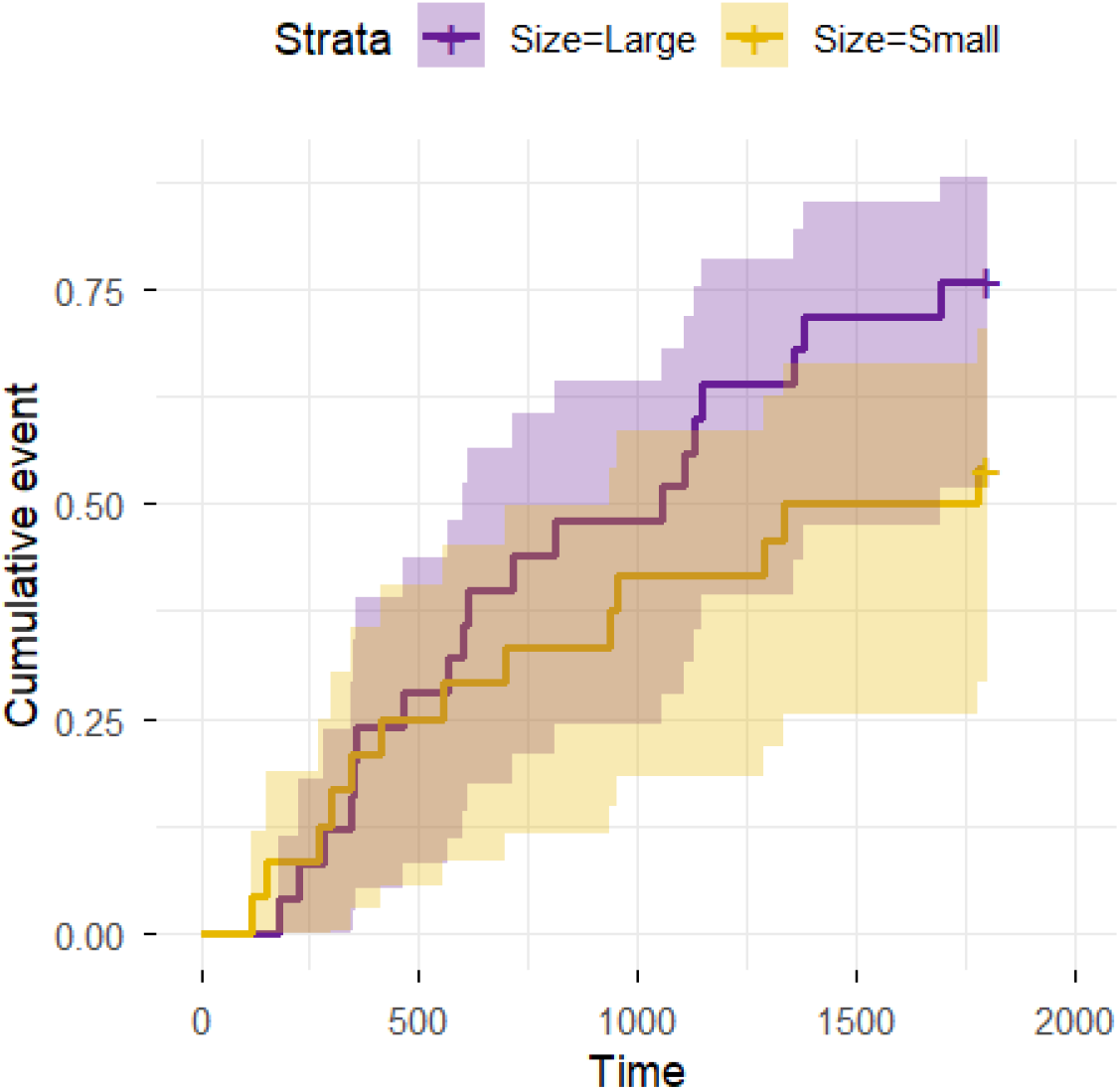
Large individuals showed a trend towards outperforming small individuals (regardless of sex) on a detour test. Cumulative event plot showing the proportion of individuals that had correctly solved the detour test at a given time within the trials, truncated at 1800 seconds when the trials were suspended. Large individuals shown in dark purple (with approximately 75% of these individuals solving the task by the end of the trial), small individuals shown in yellow (with approximately 50% solving the task by the end of the trial). Note that though the data are displayed categorically here for visualization purposes, a continuous measure of body size (thorax width) was used for the statistical model for this assay.

### Brain Gene Expression Analysis

#### Differentially Expressed Genes (DEGs)

We identified 147 genes that were differentially expressed as a function of sex, body size, cognitive performance, or that had interaction effects between these factors, as well as many genes with multiple effects. Figure 4a shows how these DEGs were distributed between categories. Supplemental Table 1 displays a similar table, but with gene names, annotated proteins, and sequences listed.

**Figure 4:**
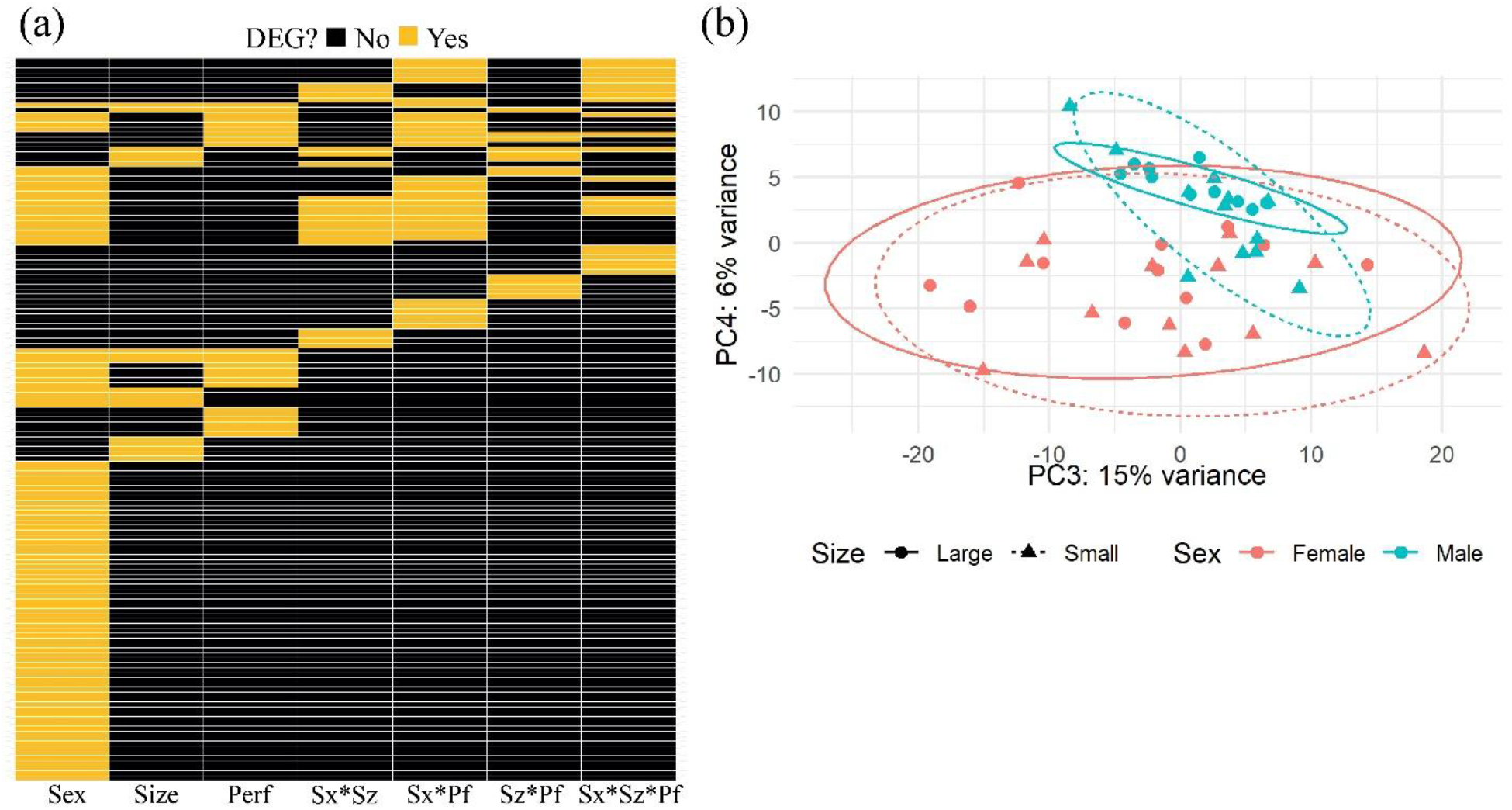
a) *Onthophagus taurus* brains show 147 genes that are differentially expressed as a function of sex, body size, cognitive performance, their interactions, or combinations of these factors. Heatmap showing the number of genes that are differentially expressed as a function of each category, in columns from left to right, of sex, body size, cognitive performance, sex*size interaction, sex*cognitive performance interaction, body size*cognitive performance interaction, or the three-way sex*body size*cognitive performance interaction. Each row represents one gene. Yellow squares indicate that that gene is differentially expressed as a function of that category, while black squares indicate that that gene is not differentially expressed as a function of that category. b) Males tended to cluster much more tightly than females for brain gene expression patterns. Principal component analysis plot showing PC3 and PC4, using data from the top 500 most variable genes. Females displayed in red, males displayed in blue, large individuals displayed with circles and solid lines, small individuals displayed in triangles with dashed lines.

The largest number of DEGs were differentially expressed between the sexes (98), including genes with known sex-specific functions, such as *vitellogenin* (a key regulator of insect oogenesis) and the insect sex determination factor, *doublesex* (a key regular of both sex- and morph-specific development in horned beetles) (Engelmann, 1979; Hildreth, 1965; Kijimoto et al., 2012). That these genes are differentially expressed in brain tissue of adults demonstrates their far-reaching consequences beyond the developmental and organizational roles that these genes are typically associated with. Eighteen genes were differentially expressed as a function of body size. This group included genes associated with regulation of gene expression, such as *Nuclear receptor coactivator 1* and a likely RNA-binding protein; these may represent candidate regulators underlying the nutrition-dependent and body size-associated male polyphenism. Twenty-three genes were differentially expressed as a function of cognitive performance. This list of genes provides a set of intriguing candidates for future functional manipulation to experimentally assess their role in cognition, including: genes associated with neurobiological function, such as a sensory neuron membrane protein, an odorant receptor, and a Kelch-like protein that is hypothesized to support neuronal development; as well as genes associated with motor performance, such as troponin. Fifty-nine genes showed at least one two-way or three-way interaction effect, and many genes were differentially expressed as a function of multiple categories. These genes highlight the complexity of the genetic machinery that is active inside the brain at a given time. Additionally, these multi-factorial and interaction-effect-laden genes could offer candidates for pathways that integrate cognitive variation with the organismal-level traits that are associated with it.

#### Overall patterns

Beyond DEGs, to consider brain gene expression as a whole, we performed a Principal Components Analysis (PCA). We found intriguing patterns associated with our independent variables of sex and body size. The first two PCs largely represented unique individual variation for each beetle that did not map onto our groups of interest. When we looked at the 3^rd^ and 4^th^ PCs, we found that males (especially major males) tended to cluster together much more tightly than females (Fig. 4b). Or to put it another way, male brain gene expression tended to be more uniform both within as well as among morphs, while female brain gene expression was more variable—at least within early principal components with relatively high explanatory power.

## DISCUSSION

In this study, we used the bull-headed dung beetle to assess the relationship between functional traits, cognition, and brain gene expression. We found that ecologically relevant life history traits such a sex and early life nutrition are associated with differences in cognitive performance. These effects are cognitive-domain-specific. And moreover, these differences are reflected in the baseline brain gene expression patterns of the individuals under study. Specifically, we found that sex was the strongest predictor of performance on a spatial cognition assay. Conversely, a continuous measure of body size—which also serves as a proxy for early-life nutrition—was the best predictor of performance on a detour task. In contrast, the categorical major/minor male diphenism was not associated with cognitive performance on any of our assays. Sex showed the most wide-ranging effects in our brain gene expression data with the highest number of DEGs, but we also identified key DEGs associated with body size and cognitive performance. Furthermore, we identified many genes that were responsive to more than one of our biological factors of interest or that showed interaction effects. And finally, we found that the brain gene expression signatures of males were much more similar to one another than for females. Intriguingly, this effect was strongest for major males, which were in the highest-performing groups on both assays (“males” for the spatial cognition assays and “large individuals” for the detour task).

We had hypothesized that our functional groups would perform similarly on average on the spatial cognition assay, as all beetles must navigate the complex labyrinth of tunnels that they excavate underneath the dung pads. We instead found that sex was a significant predictor, with males outperforming females. This result could suggest that there is stronger selective pressure for males to have a thorough understanding of tunnel architecture during reproductive contests (both for majors who are defending tunnels and minors who are attempting to sneak past the majors), compared to females who mainly work in a single tunnel at a time while constructing brood balls. Alternatively, females might not have been failing to learn, but rather their at-chance-levels of performance in our assay could suggest that they were showing a greater tendency to actively explore the tunnel ecosystem. Brood parasitism is documented to be a common and rewarding strategy for females, particularly under stressful conditions, so females may benefit from continuously exploring all available tunnel options (Moczek & Cochrane, 2006). It is also possible that females would have shown higher performance in a different learning context, such as if we had chosen to motivate them with opportunities to construct brood balls rather than with a food reward.

We had also hypothesized that minor males (who routinely switch between different reproductive tactics of fighting and sneaking) would perform the best out of our functional groups on the detour test. Instead, we found that our statistical model performed better when we used a continuous measure of body size (which is strongly associated with early life nutrition) rather than the binary major/minor diphenism; and furthermore, when we did so, that large size (rather than the small size of minor males) emerged as the best predictor of performance on this assay, regardless of sex. The enhanced performance of large individuals might be explained by body-size-dependent differences in nutritional buffering capacity. Cognition in general is believed to be energetically expensive, and some of the cognitive skills that are typically associated with a detour task (such as inhibitory control) are especially sensitive to nutritional state (Bartholdy et al., 2016; Christie & Schrater, 2015). It is therefore possible that our larger individuals possessed greater long-term nutritional reserves and were consequently better able to maintain this cognitive capacity even under mild food deprivation.

Our brain gene expression results provide avenues to better understand these cognitive findings and provide fodder for future mechanistic studies. For example, sex was an important predictor of cognitive performance. We found 33 genes with a statistically significant interaction effect between sex and cognitive performance, meaning that the expression levels of those genes relative to one factor *depended on* the state of the other factor. For example, the gene that codes for the protein “General odorant-binding protein 56d” was one gene with an interaction effect between sex and cognitive performance (Fig. 5a). Odorant binding proteins in insects typically operate within the antennae to facilitate odorant detection, but they have also been observed to act within the central nervous system in the context of maintaining lipid homeostasis (Yin et al., 2025). High-performing individuals consistently showed lower expression levels of this gene than low-performing individuals, but these dynamics were influenced by sex; males more generally showed lower expression than females. We can thus map these expression patterns onto our cognitive results: i) males generally performed better on the spatial cognition assay than females; ii) males, the higher-performing sex, showed lower expression of the gene that codes for General odorant-binding protein 56d than females; iii) the females that *did* show higher cognitive performance showed lower expression than low-performing females of this gene. Therefore, lower expression of the gene that codes for General odorant-binding protein 56d seems to be generally associated with better performance on the spatial cognition assay across two dimensions—lower expression generally in the sex that performed better, and low expression within sexes in the individuals that outperformed their peers. This gene may therefore be important for spatial cognition in particular and would be a promising candidate for future knockdown manipulations.

**Figure 5:**
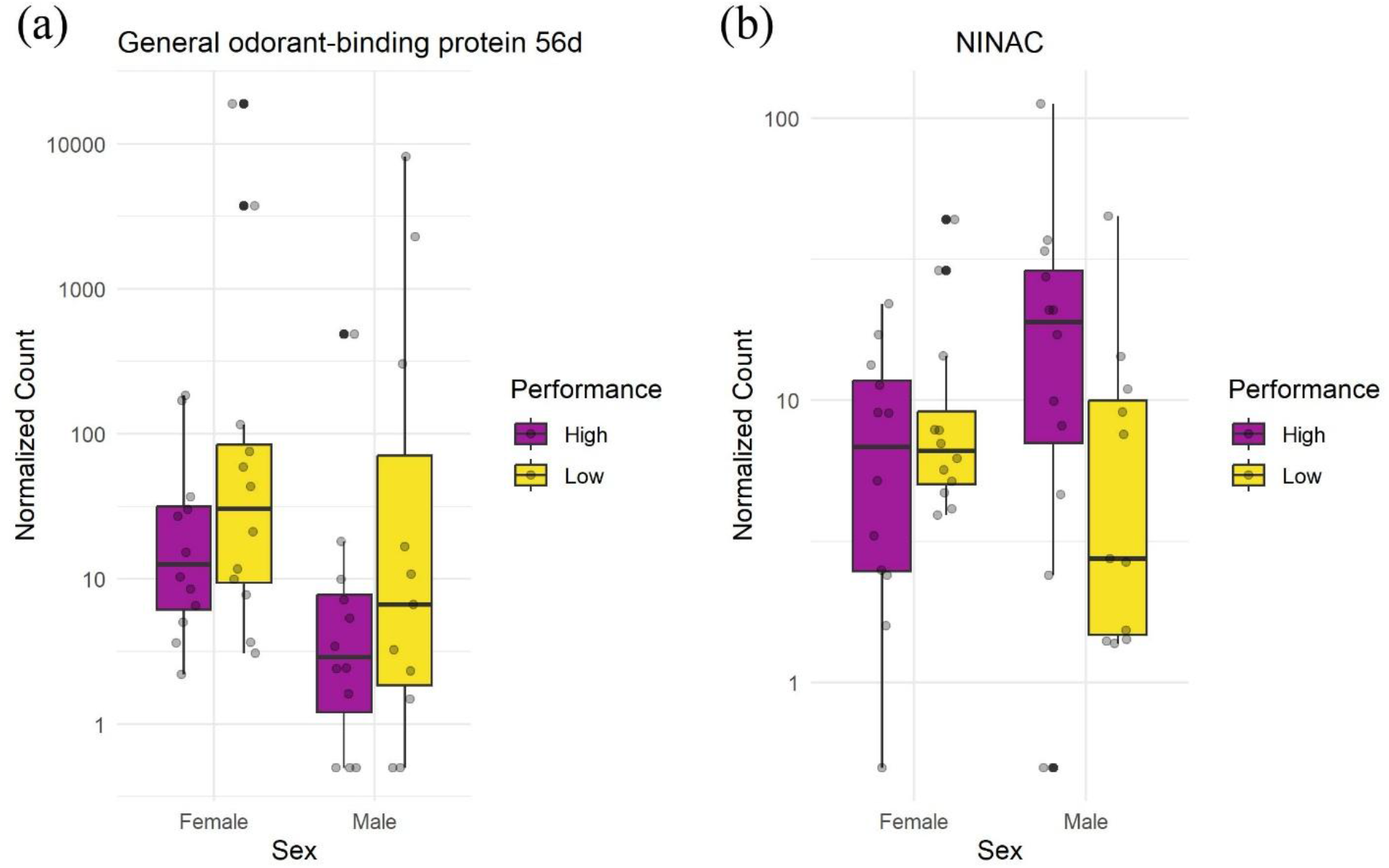
Two examples of genes that showed statistically significant interaction effects between sex and a composite measure of cognitive performance. Both plots show females on the left and males on the right, high-performing individuals in purple and low-performing individuals in yellow. Note the use of a log10 scale on the y-axis for readability. a) Boxplots showing normalized expression levels of the gene “General odorant-binding protein 56d”; b) boxplots showing normalized expression levels of the gene “Neither inactivation nor afterpotential protein C” (NINAC).

Another example of these dynamics can be seen with the gene that codes for “Neither inactivation nor afterpotential protein C” (ninaC). Though this gene has been best studied for its role in photoreceptor cells, it has been documented to be expressed in the brain in ants, where it shows differential expression levels between workers and gynes (Qiu et al., 2018). In our data, we found that the gene that codes for ninaC showed similar expression levels in females regardless of performance, while in males it showed higher expression in high-performing individuals than low-performing individuals (Fig. 5b). Some genes may therefore be more important for particular subgroups than others, and different groups may rely on different neurobiological substrates to achieve similar behavioral outcomes.

These principles can be applied to our other groups as well. For instance, we found that body size was associated with performance on the detour task. We found 13 genes with statistically significant interaction effects between body size and cognitive performance. One of these was the gene that codes for the protein “Sodium-independent sulfate anion transporter” (SLC26A11). This gene is highly expressed in brain tissue and is important for both sulfate and chloride exchange, with implications for neurobiological pathology (Kuhn et al., 2026). We found that the gene that codes for SLC26A11 is particularly highly expressed in one of our subgroups: high-performing, large individuals (Fig. S2). As large individuals generally outperformed small individuals on the detour task specifically, this gene may be important for the cognitive skills involved in solving this task. Many other genes with significant interaction effects were identified in this study that could each be examined for their significance in linking our traits of interest (Table S1).

When examining broader brain gene expression patterns, we additionally found that males showed much more tightly clustered gene expression patterns than females. It might at first appear counterintuitive that the sex with two distinct behavioral types (major “fighters” and minor “sneakers”) would show *more* narrow expression patterns. On the contrary, it is possible that the specialization required to produce these distinct phenotypes acts as a constraint that limits what expression patterns are viable. Females, on the other hand, display a range of complex behaviors (context-dependent parental care decisions, brood ball construction, brood parasitism, intra- and intersexual aggression, foraging, tunnel excavation, etc.) and do not need to be herded into one of two neat behavioral “bins.” This freedom could allow more diverse brain gene expression states to exist without causing detrimental effects on the behaviors they express.

It is important to note that our RNAseq results measured brain gene expression at baseline rather than in close temporal proximity to the completion of any of our assays. We chose to measure it this way because no cognitive assessments had ever been executed in this species of dung beetle prior, and we therefore did not go into this experiment with a particular expectation about which cognition test would be the most salient. This approach also allowed us to capture the substrate of expression that a beetle brain would be working with when first confronted with a novel situation that required cognitive input. It is conceivable based on previous research that our differential expression results would have been stronger if we had measured brain gene expression within minutes to hours after completion of one of the assays (Cullinan et al., 1995). Future work could explicitly measure how these groups differ in the immediate change in expression following a learning event. And finally, our analysis identified several intriguing candidate genes that could be targets for future functional manipulation (such as RNA*interference*, which is well-established in *Onthophagine* beetles) to more carefully probe their involvement in cognitive performance.

Overall, this research illustrates the importance of considering multiple ecologically relevant traits and multiple cognitive domains when assessing individual variation in cognition. This work additionally highlights the role of brain gene expression as a mechanism by which these traits may can be integrated across a whole organism.

## Acknowledgements

This work was supported by funding from the National Science Foundation [Grant no. 1901680, awarded to APM; Grant no. 2508006, Postdoctoral Research Fellowship in Biology, awarded to RRW]; and the National Institutes of Health [Common Themes in Reproductive Diversity training fellowship, awarded to RRW]. We would additionally like to extend sincere thanks to Kip and Whitney Schlegl of Marble Hill Farm for allowing us to collect the cow dung used to feed and reward the beetles in this experiment; The Van Nieuwenhze lab, especially Dr. Zachary Taylor, for allowing us to use their lyophilizer; Dr. Erica Nadolski for the use of her beetle photos, as well as her assistance with training RRW; our beetle colony care assistants, Isabel Manley and Eve Pieri; and the Indiana University Center for Genomics and Bioinformatics for their sequencing expertise.

## Supplemental Materials

Table S1: [Separate document]

**Figure S1:**
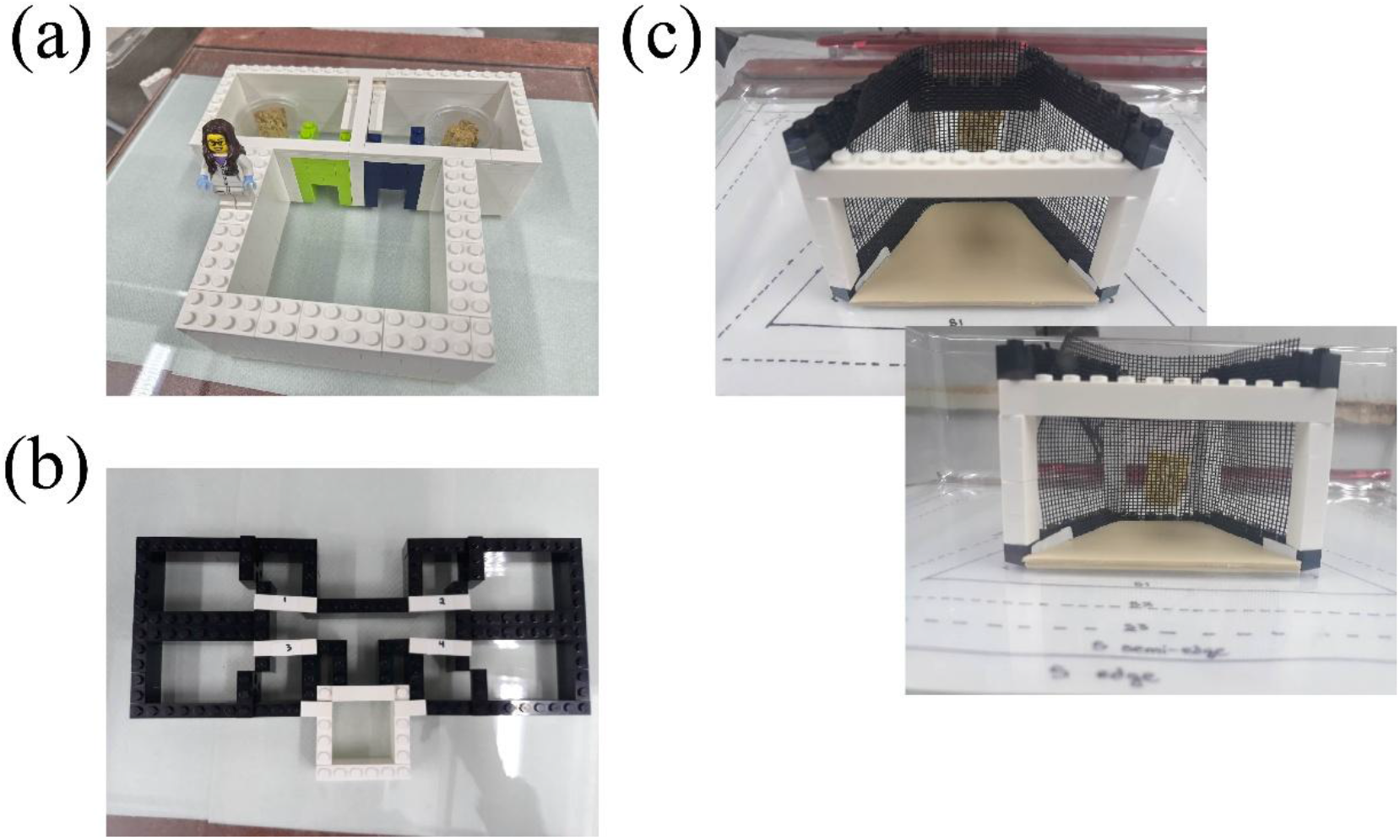
LEGO™ arenas used in this experiment; a) associative learning arena; b) spatial cognition arena; c) detour test apparatus.

**Figure S2:**
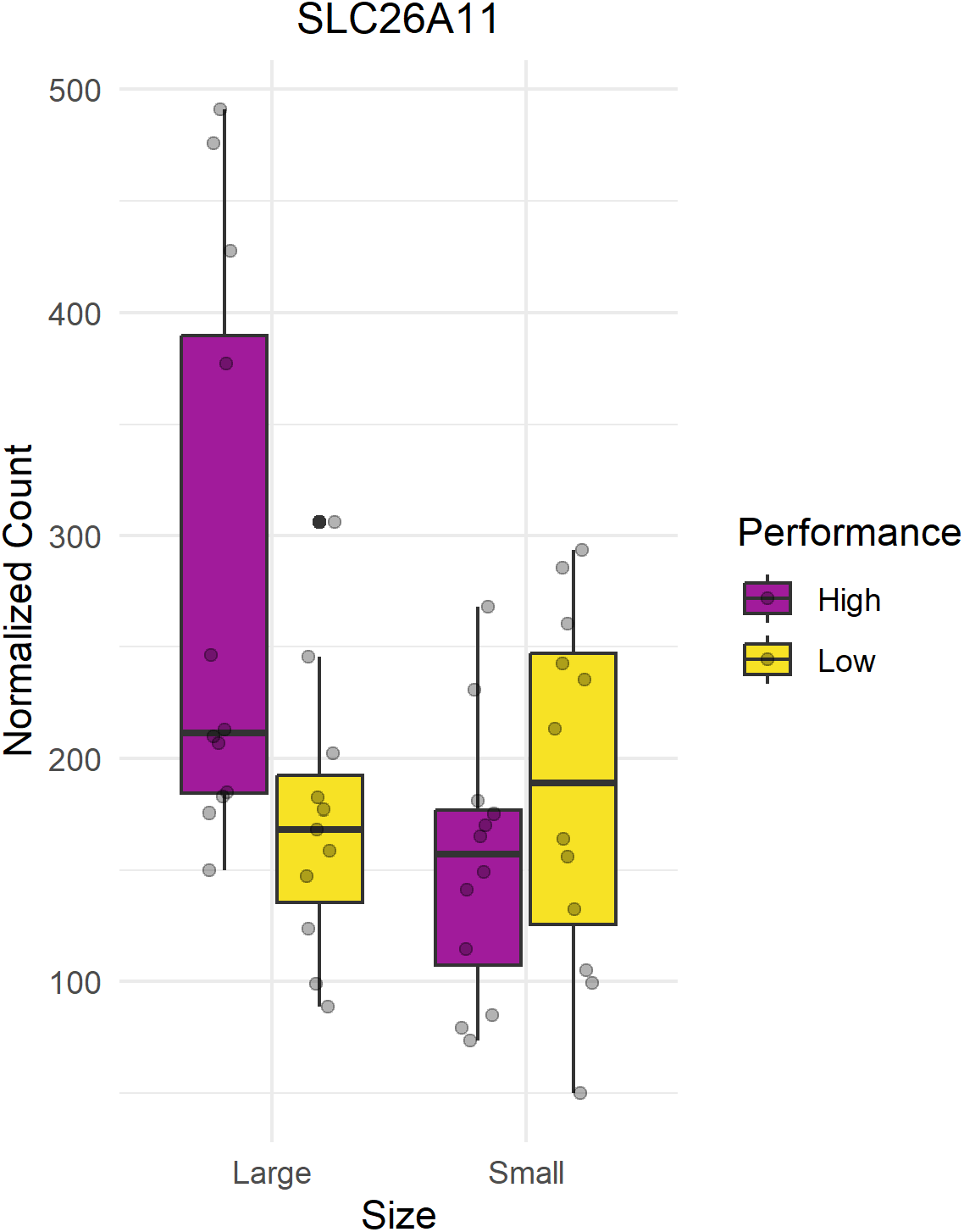
The gene that codes for the protein “Sodium-independent sulfate anion transporter” showed a statistically significant interaction effect between body size and cognitive performance. Boxplots show large individuals on the left and small individuals on the right, high-performing individuals in purple and low-performing individuals in yellow.

